# Culture-free single-cell transcriptomics shows short-term functional and associative stability in non-model microeukaryotes under shading stress

**DOI:** 10.64898/2026.09.07.749927

**Authors:** Aditya Jeevannavar, Javier Florenza, Stefan Bertilsson, Manu Tamminen

**Author notes:** Corresponding author: Manu Tamminen (; Department of Biology, University of Turku, Vesilinnantie 5, 20500 Turku, Finland).

## Abstract

Microeukaryotes are abundant, diverse, and morphologically and functionally complex. They represent one of the largest groups of primary producers and are involved in major ecosystem functions such as nutrient transformations and maintenance of food webs. Although ∼74,000 species of microeukaryotes are estimated to exist, very few are available in culture and only ∼3000 are represented by reference genomes and transcriptomes. This limited representation significantly hinders the study of microeukaryotes in their natural environments. Single-cell transcriptomics with sequencing of full-length transcripts has potential to bypass this limitation. In this study, we employ Smart-seq3xpress for taxonomic identification and functional characterization of non-model mixotrophic freshwater microeukaryotes in a culture-free and reference-free manner, while also enabling inferences to be made for their prokaryotic associations. Computational analyses revealed 22 distinct microeukaryote taxa upon sequencing of 1520 randomly sampled cells, with assemblies of 12 good quality *de novo* partial transcriptomes, followed by functional annotation using sequence and structure homology. Gene expression analysis of the transcriptome for the most abundant microeukaryote in the sample, *Rhodomonas*, revealed a transcriptomic dip a few hours after experimental shading, followed by a gradual recovery in the next few days. Compared to samples in illuminated conditions, there was widespread downregulation of photosynthesis-, lysosome-, and carbon metabolism-related pathways in the shade, but the prevailing associations with prokaryotic lineages was unaffected by the light conditions. The different microeukaryotes, all present in the same environmental sample, featured distinct prokaryotic associates that were consistent across illumination conditions. This reference-free and culture-free study of 22 different microeukaryotic taxa sets the stage for high-throughput single cell transcriptomics to study microeukaryotic diversity in complex natural ecosystems, for their population-level taxonomic identity, sub-population-level transcriptomic profiles and metabolic states, and individual-level prokaryotic associations.

## Introduction

Microeukaryotes are abundant (estimated to be of the order of 10^27^)^1^ and represent the majority of the eukaryotic tree of life^2,3^. They are diverse in terms of trophic behaviour (presenting phototrophic, phagotrophic, and mixotrophic modes)^4^, morphology (bearing flagella, cilia, pseudopodia etc)^5^, and molecular traits (possessing complex transcriptional mechanisms like twintrons and 4-or 5-letter codons)^6^. However, detailed genomic, morphological, ecological, and evolutionary study of these microeukaryotes has been hampered by the difficulty of cultivating most of this diversity^7^.

Microeukaryotes engage in various ecosystem functions including large-scale primary production, variety of symbioses, and operating multiple roles in the food web^8^. Aside from taxonomically identifying the microeukaryotes and phylogenetically classifying them, there is great interest in deciphering their functional diversity across populations and subpopulations. Due to the lack of microeukaryote sequences in public databases, intensive manual curation for functional annotation, metabolic mapping, and identification of orthologs and paralogs is necessary^9^. This manual curation is possible when studying a single organism but becomes increasingly difficult when scaling up to studying dozens or hundreds of organisms simultaneously to cover the full diversity in natural ecosystems.

Biotic interactions are critical for the ecosystem functioning of microeukaryotes. Microeukaryotes generally have prokaryotic ecto-and endosymbionts^10^. Some microeukaryotes form symbiotic relations with corals, flatworms, anemones, and mollusks^11^. Many of the protists prey on prokaryotes and may contain prey-associated sequences in sampled transcriptomes^10^. Some are also known to impose strong selection on the associated prokaryotic community and forming mutualistic metabolic relationships^8,12^. After taxonomic identification and functional characterization, interaction with prokaryotes may shed further light on their biology and function in the ecosystem. Although there are difficulties in separating out the prokaryotic “contaminant” sequences^10^ the opportunity to study inter-species interaction using transcriptomic data has so far not been widely recognized, and consequently no widely-used methods to do so exist.

Beyond phylogenomic analyses, the study of even those microeukaryotes that we have managed to culture or whose genomes we have managed to sequence is obscured by their invariably complicated transcriptional mechanisms, to the extent that one might not be able to infer the mature mRNA transcript sequence, and consequently describe the translated protein and its function from the mere gene sequence^6^. Large international efforts, such as the Marine Microbial Eukaryotic Transcriptome Sequencing Project (MMETSP)^13^ and the Protist 10,000 Genomes Project (P10K)^14^, have been launched to create a collection of transcriptomic and genomic reference datasets. However, as mentioned above, genomic references might be ineffective until we can confidently predict transcripts, and consequently their inferred functions, from genome sequence data. Different problems arise while using transcriptomic references, in terms of culturableness and coverage. Culture-based reference transcriptomes, like those from MMETSP, cover much of the transcriptional capacity of microeukaryotes but require that the microeukaryotes are available in culture. In contrast, single-amplified reference transcriptomes do not rely on culturing but cover only the sampling condition dependent partial transcriptional capacity.

Although 74,000 species of protists are estimated to exist, only ∼3,000 reference genomes and transcriptomes are so far available for this group^14,15^. Considering that even references from different species in the same genus yield scant read mapping, these currently available genomic and transcriptomic references are scarce. Most protists are not available in culture either. Thus, reference-free single-cell transcriptomics currently appears to be the most rational and feasible way to study the diversity of these microeukaryotes^16–18^.

Previous single-cell transcriptomics studies on microeukaryotes have focused on species in culture^17,19^, model microeukaryotes like malarial parasites^20,21^ and yeasts^22,23^, manual picking of individual non-model cells^24–29^, or solely on phylogenomic analyses^30,31^. Although all these studies successfully characterized gene expression profiles in microeukaryotes, they bypassed the complexities of natural microeukaryote systems such as unculturability, high phylogenetic and size diversity, and low abundance^32^. Two other studies focussing on microeukaryotes’ single-cell gene expression stand out due to their potential for use in natural communities, one using Smart-seq2 on *Giardia intestinalis* cells^33^ (the method used in our previous proof-of-concept with *Ochromonas triangulata cells*^34^ and the non-UMI precursor method to the Smart-seq3xpress used in this study) and one using MASC-seq with *Tetrahymena thermophila*, *Phaeodactylum tricornutum*, and *Heterocapsa* sp cells^35^. They both apply the methods using microeukaryotes with reference genomes or transcriptomes obtained outside of the single-cell transcriptomic sequencing, but showcase the potential for culture-free microeukaryote expression study.

Full-transcript sequencing methods, such as Smart-seq3xpress, have the potential to enable functional characterization of microeukaryotes lacking reference genomes and transcriptomes^36^. The use of Unique Molecular Identifiers (UMIs) enables accurate transcript counting. Although oligo-dT primers are used in the protocol to enrich for mRNAs, the massive amount of rRNA in the cells typically leads to substantial carryover of rRNA into the final sequenced RNA pool. This carryover rRNA, typically discarded as non-target sequences, can be used for taxonomic identification of the microeukaryote as well as their ecto-and endosymbionts manifested at the single-cell level.

In a previous study, we demonstrated the capacity of single-cell transcriptomics for taxonomic identification, functional characterization, and prokaryotic association inferences for a non-axenic non-model mixotrophic microeukaryote. In this follow-up study, we expand this methodology to the mixotrophic subset of natural microeukaryotic communities in a freshwater lake. A natural lake water sample was subjected to two distinct illumination conditions and sampled at various time intervals. Mixotrophic cells were isolated using fluorescence-activated cell sorting (FACS) and single-cell transcriptomic libraries were prepared using Smart-seq3xpress. Computational analyses of the full-length transcripts were used for identification of the microeukaryotes and their prokaryotic associates, as well as characterization of their gene-inferred functions demonstrating the possibility of reference-free and culture-free study of uncultivated microeukaryotes using single-cell transcriptomics.

## Results

### Expression in *de novo* assembled transcriptomes clusters by taxa

Natural samples from oligotrophic, clear water lake Långsjön (Sweden, 60° 1’ 58.5’’ N, 17° 34’ 47.1’’ E) were acclimatised for 3 days, subjected to experimental shading, sampled at different time intervals, and sorted to select for mixotrophic activity (Fig 1). The eukaryote abundance remained comparable across the treatments except for the decline in the shaded flasks at the 48h timepoint (Fig 2A). Bacterial abundance also remained comparable across treatments except for the decline in the shaded flasks at the 24h timepoint (Fig 2B).

**Figure 1:**
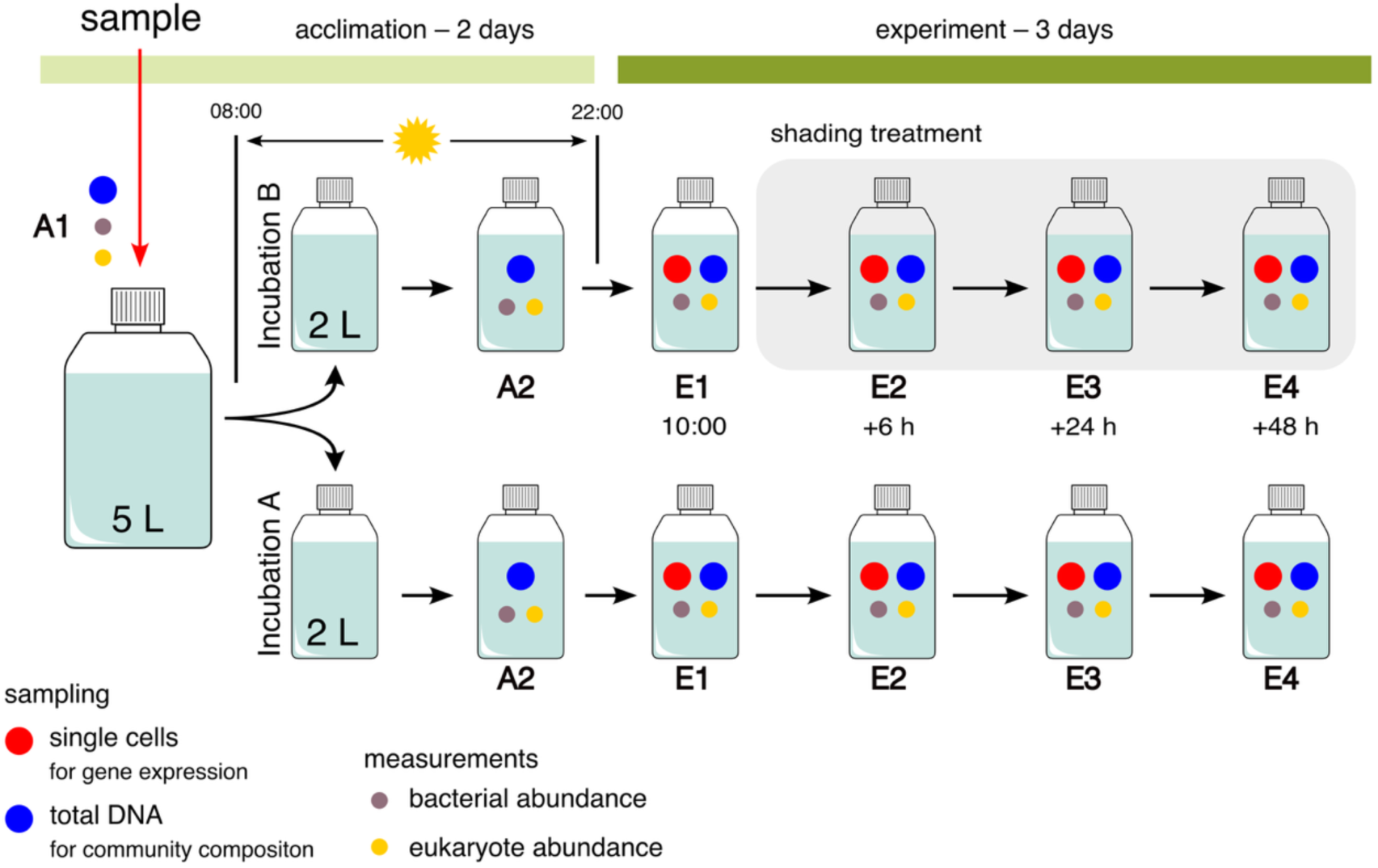
Schematic of the experimental design: Samples from Laken Långsjön were taken on October 8^th^, 2024, and acclimatized in a 5L flask for 2 days in a 14-hour photoperiod. Subsequently, they were divided into two batches receiving different treatments: illuminated treatment (14h photoperiod at 80 μmol s^-1^ m^-2^ irradiance) and shaded treatment (14h photoperiod at 6 μmol s^-1^ m^-2^ irradiance). Samples were taken 0, 6, 24, and 48 hours after the onset of treatment conditions.

**Figure 2:**
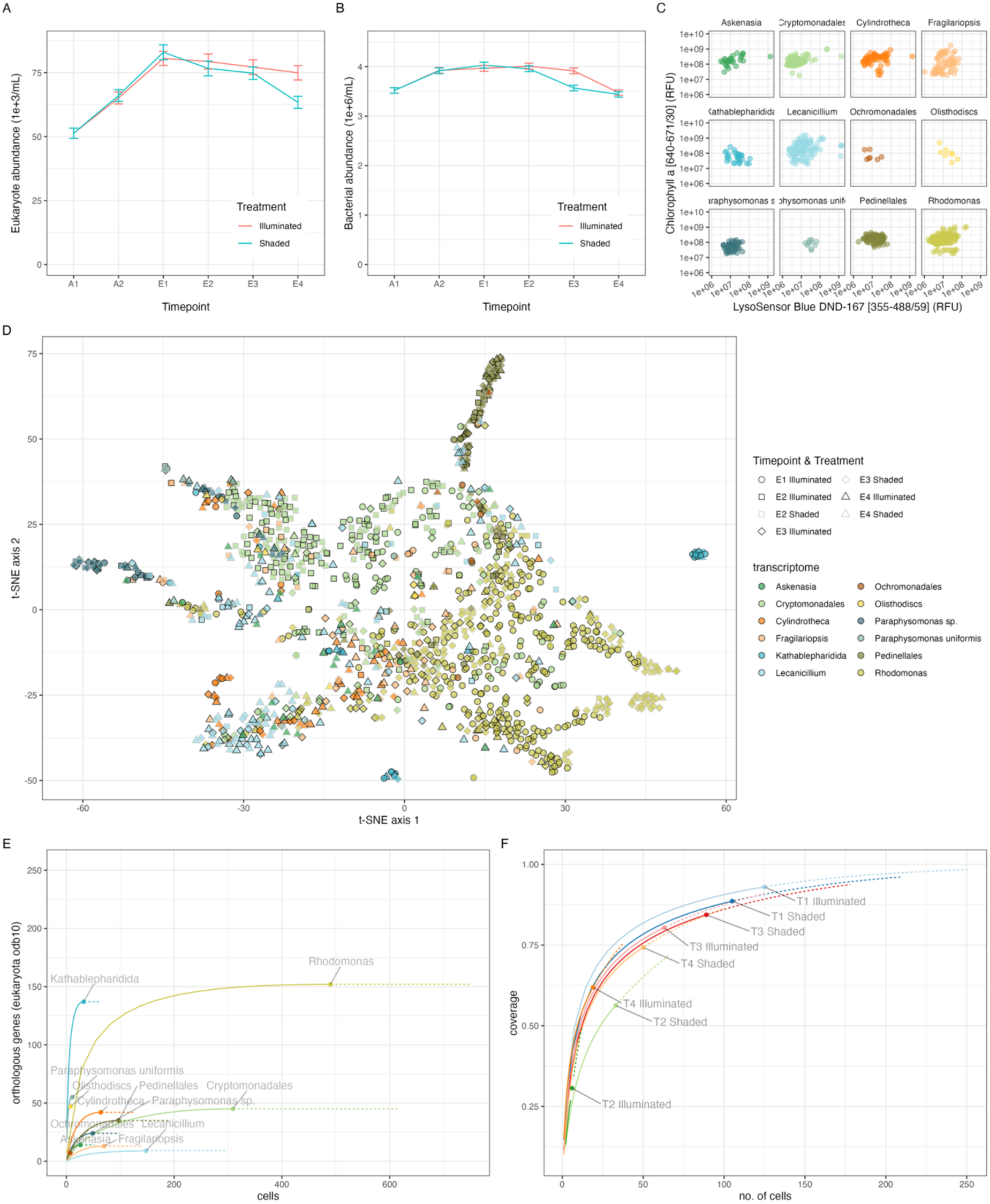
Sampling, sorting, and clustering the microeukaryotes: (A), (B) Abundance (ml^-1^) of eukaryotes and prokaryotes in the acclimatizing and experiment flasks. (C) Flow cytograms of FACS-sorted cells for each of the 12 largest transcriptomes in the study. (D) t-SNE visualization of the gene expression based on genes expressed in the 12 largest transcriptomes in the study. Each dot represents a cell whose colour is based on rRNA-based taxonomic assignment and outline is based on treatment and timepoint. (E) Partial transcriptomes’ accumulation plot for the 12 transcriptomes’ samples. Transcripts of the eukaryota orthoDB genes (n = 255) expressed in and recovered from the cells were considered. (F) Partial transcriptome coverage plot for each sampling cluster of the *Rhodomonas* transcriptome. For (E) and (F), the dots represent sample size, solid lines represent sample rarefaction, and dotted lines represent extrapolation based on rarefaction model for each group.

1,520 cells were sequenced (190 per time interval and treatment), of which 46 were excluded due to too few UMIs. On average, 859,794 read pairs were recovered per cell, of which 747,269 were UMI-end reads. After trimming and contaminant sequence filtering, a median of 2,192 unique UMIs mapped to the assembled transcriptomes. Among the 1,474 cells, 22 distinct taxa were detected based on non-specific recovery of ribosomal RNA (Supplementary table 1), and 22 distinct transcriptomes were subsequently assembled *de novo*. 18 of the 22 microeukaryotic identities were recovered using the rRNA transcript-based taxonomy assignment. The microeukaryotic identities of 99.96 % of these cells, at family level, were recovered from the 18S rRNA amplicon sequencing of the samples.

The largest transcriptomic cluster, *Rhodomonas sp.*, consisted of 490 cells and 227,638 distinct transcripts, while the smallest (*Haslea, Paulinella,* and *Telonemia)* consisted of 1 cell each (75, 88, and 3,962 transcripts respectively, Supplementary table 2). While the FACS signals for photosynthetic activity and acidic vacuole presence mostly selected for putative mixotrophs, occasional putative heterotrophs like katablepharids also passed the filters, likely due to the ingestion of photosynthetic bacteria (Fig 2C).

The 12 biggest annotated transcriptomes (covering 88.7% of the 1,474 cells) had a median of 25,423 transcripts in the *de novo* assemblies, with a median of 5,920 annotated transcripts and 98 of these expressed across the 12 groups. Visualization and clustering based on these commonly expressed transcripts show that the gene expression clusters coincide with the taxonomic origin of the transcriptomes (Fig 2D). Hierarchical clustering using Bray-Curtis dissimilarity (with tree cut at half the maximum height) resulted in clusters corresponding to the taxonomic origin of the transcriptomes (Rand index = 0.77).

Rarefaction and extrapolation of the expression of orthologous genes in the eukaryota orthoDB list (containing 255 genes) revealed that Rhodomonas (152 out of 255 genes) and Kathablepharidida (137 out of 255 genes) expressed the largest number of these orthologous genes while having vastly different number of recovered cells, i.e. 490 and 32 respectively (Fig 2E). However, subsetting the cells based on treatment and sampling time showed that the coverage of the *Rhodomonas* transcriptome depended heavily on the number of cells recovered (Fig 2F).

### Microeukaryote-prokaryote association are microeukaryote species-specific and unaffected by light conditions

Based on the non-specific recovery of bacterial rRNA, *Endomicrobiaceae* (with Alveolates and Stramenopiles) and *Leptolyngbyaceae* (with cryptophytes) were the two most common prokaryotic associates of the microeukaryotes, followed by *Microcystaceae* (with *Lecanicillium* and *Paraphysomonas* cells) (Fig 3A, 3B). The presence of *Leptolyngbyaceae* and 76.34% of other prokaryotic families forming the associations was confirmed from bulk DNA by 16S rRNA amplicon sequencing, with the exception of Endomicrobiaceae which was not observed in the bulk samples.

**Figure 3:**
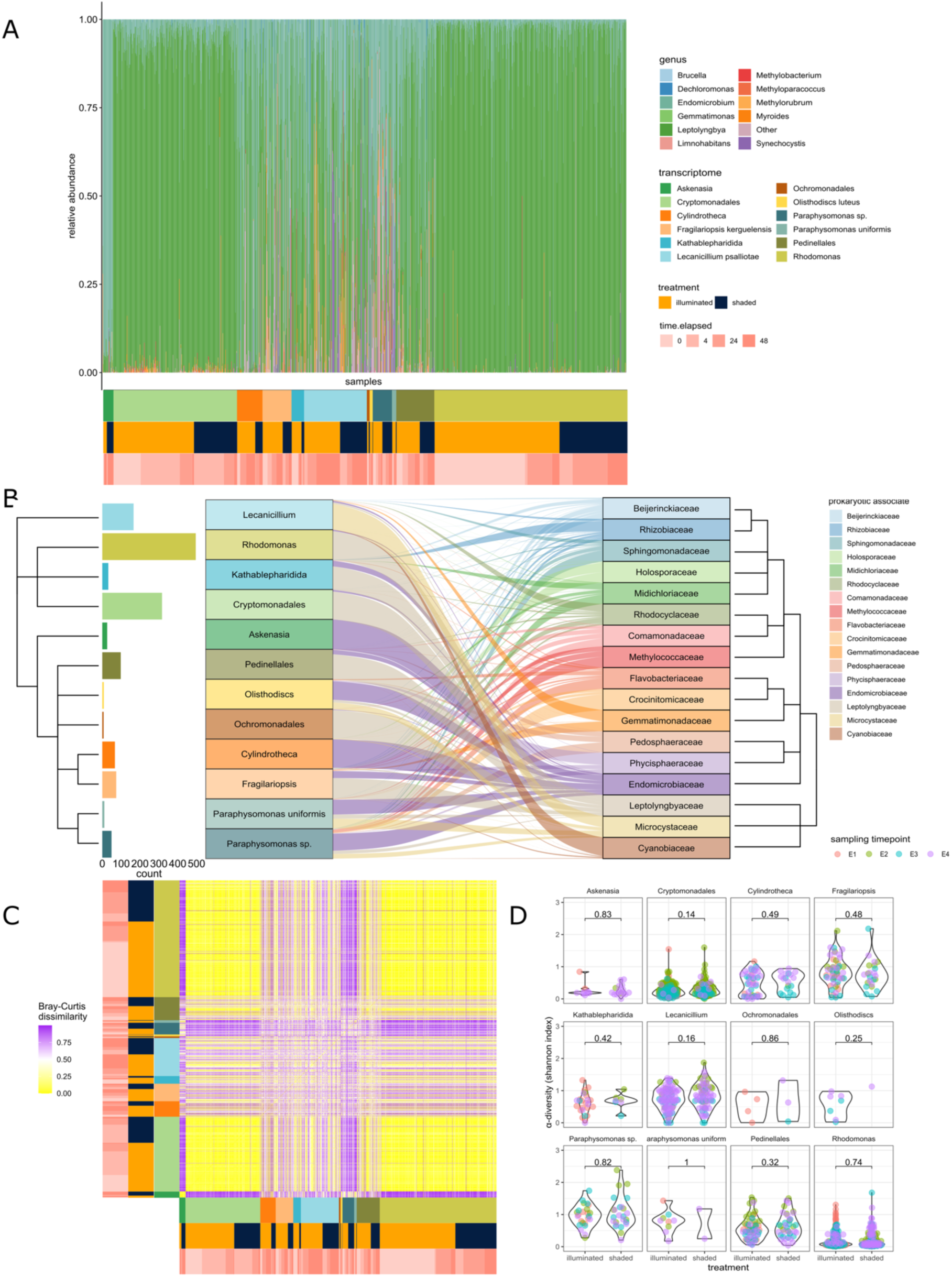
Associated prokaryota: (A) Relative abundance of the most abundant prokaryotic genera associated with the different microeukaryotes (indicated at the bottom of the x-axis) in different treatments (orange-blue colour bar under x-axis) along different sampling timepoints (peach colour bar under x-axis). (B) Alluvial plot of each of the microeukaryote’s associations with prokaryotic families. The thickness of the flows represents relative abundance of the prokaryotic associated, normalized on both ends of the flow. The bar chart on the left of the alluvial plot represents the number of cells sampled from each microeukaryote transcriptome. The phylogenetic trees were generated from NCBI Taxonomy using the NCBI Common Tree tool. (C) Bray-Curtis dissimilarity of each sample’s associated prokaryota with the rest of the samples with the different microeukaryotes (indicated at the bottom of the x-axis and left of y-axis) in different treatments (orange-blue colour bar) along different sampling timepoints (peach colour bar). (D) Alpha diversity of each sample’s associated prokaryota facetted by transcriptome, separated by treatment condition, and coloured by sampling time.

The community of prokaryotic associates (detected by single-cell transcriptomics, average shannon diversity = 0.65) exhibited a lower alpha diversity than the community in the bulk sample (detected by amplicon sequencing, average shannon diversity = 4.85) and the two were vastly different in their composition (p = 0.0001, F = 18.2558, PERMANOVA), indicative of only a subset of ambient prokaryota participating in symbiotic or prey association with the microeukaryotes.

The associations were observed to be specific to microeukaryote species. The prokaryotic communities were more homogenous intraspecies (average Bray-Curtis dissimilarity = 0.16±0.22) compared to interspecies (average Bray-Curtis dissimilarity = 0.43±0.33, Fig 3C). Multiple families of bacteria, like *Leptolyngbyaceae, Endomicrobiaceae, Microcystaceae, Flabovacteriaceae, Beijerinckiaceae, Comamonadaceae, Phycisphaeraceae, Gemmatimonadaceae, Rhozibiaceae, Rhodocyclaceae, Pedosphaeraceae, Crocinitomicaceae, Sphingomonadaceae, Cyanobiaceae, Holosporaceae*, were found to be differentially abundant in their associations with different microeukaryotes (Supplementary table 4).

Generally, the prokaryotic association for the different microeukaryotes did not change as a response to the treatment (Fig 3C), with the exception of *Leptolyngbyaceae, which* was found to be differentially abundant between the two treatment conditions in its association with *Lecanicillium*. Permutational ANOVA of a simple additive redundance analysis model showed that the biggest explanatory factor for the relative abundance of the prokaryotic associates was the taxonomic identity of the microeukaryote (F = 51.5499, p = 0.001), followed by sampling time (F = 9.6232, p = 0.001). The treatment did not have a significant effect on neither relative abundance nor alpha diversity of the associated prokaryotic taxa (Fig 3D).

### *Rhodomonas*, the most abundant mixotroph, dips in transcriptional activity after 6 hours of shading

Reporting of transcriptional inferences have been limited here to the most abundant mixotroph *Rhodomonas*. The *Rhodomonas* de novo transcriptome was assembled by combining reads from 491 cells, nearly a third of all sequenced cells. *Rhodomonas* cells had a median of 553,151 reads recovered and a median of 37,344 unique UMIs, of which 2,192 mapped to the assembled transcriptome. Rhodomonas cells were recovered in all treatments and timepoints, with the highest at timepoint 1, with 125 and 105 cells in incubations A and B respectively, and lowest at timepoint 2, with 6 and 33 cells in incubation A and B respectively. 400,667 contigs were assembled de novo after pooling the reads from the individual *Rhodomonas* cells of which 227,638 transcripts remained after filtering for open reading frames and longest isoforms. 68,224 of the above transcripts were annotated.

424 genes were differentially expressed between the illuminated and shaded treatments (Fig 4A, Supplementary table 3). However, excluding samples from timepoint 1 (taken immediately after onset of treatment conditions) reduced the number of differentially expressed genes to 80. Between the treatments, when considering sampling timepoints 2, 3, and 4 individually, the numbers were only 23, 20, and 18 respectively. Compared to timepoint 1 of incubation B (shaded), 275 genes were downregulated for timepoint 2 while only 4 genes were upregulated, indicating a dip in transcriptional activity 6 hours after the onset of shading. At timepoint 3, 180 genes were downregulated and 65 upregulated, indicating partial persistence of the transcriptional dip. For timepoint 4, only 85 genes were downregulated and 23 genes were upregulated, indicating a partial recovery of transcriptional activity.

**Figure 4:**
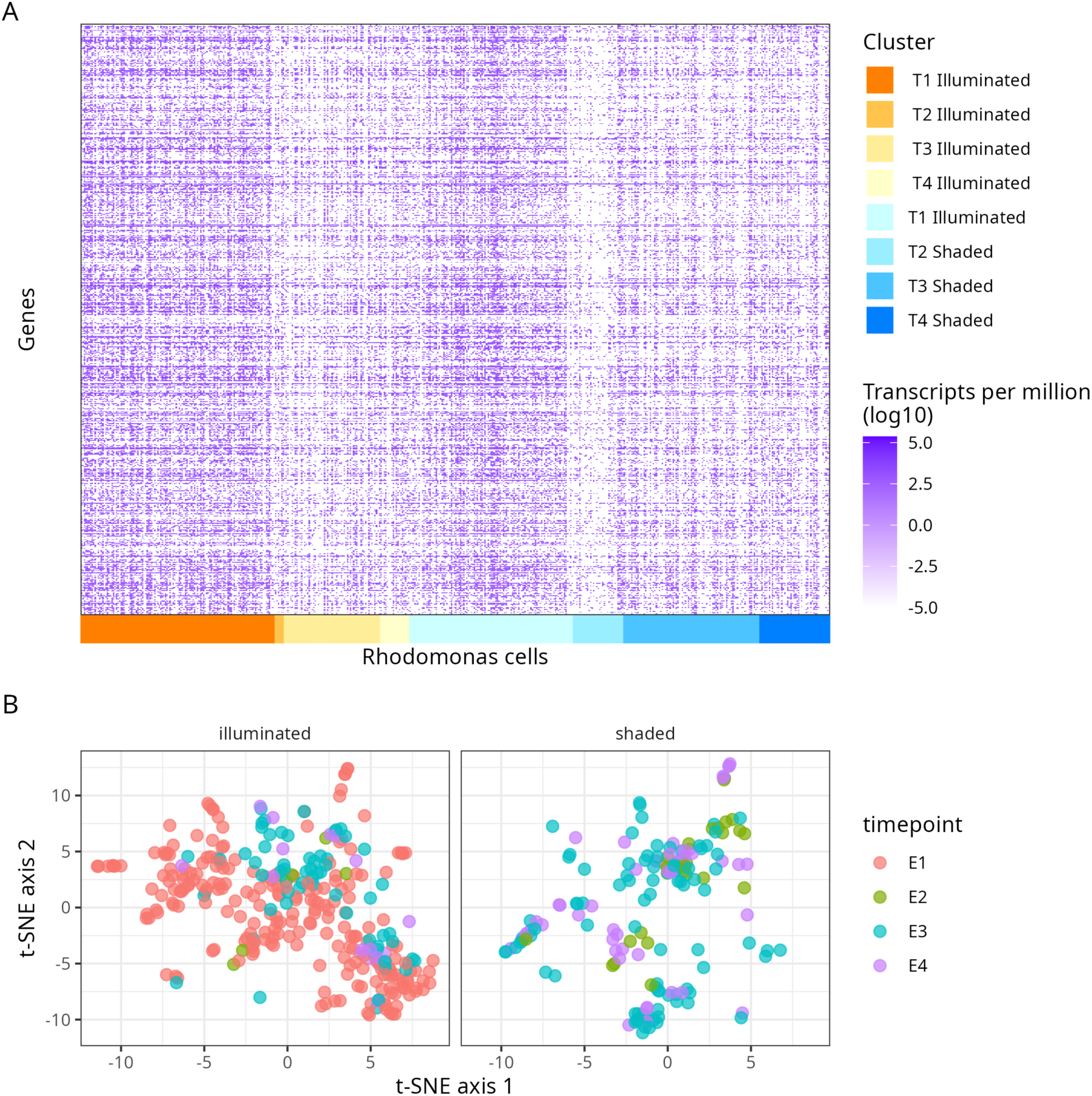
*Rhodomonas* gene expression: (A) Heatmap of genes that were differentially expressed between illuminated and shaded treatments. Gene expression of the cells has been ordered by treatment and sampling time, and the colour bar at the bottom of the x-axis represents this. Timepoint 1 of incubation B (which was shaded) was part of the illumination treatment since no effect of shading is expected to be seen instantaneously. (B) t-SNE visualization of the differentially expressed genes’ expression in *Rhodomonas*, and coloured by sampling time, and facetted by treatment, i.e., t-SNE was done with all cells together (as indicated by the same axis on both facets) and presented in two facets only for clarity.

Clustering of gene expression using shared nearest-neighbour graph and hierarchical clustering (with cosine dissimilarity) enabled the complete recovery of illuminated (Rand index = 1 for both methods) and shaded (Rand index = 0.89, 0.71 for the two methods respectively) clusters at timepoint 2, however they could not be separated from each other (Rand index = -1.09, -1.32 for the two methods respectively). Resolution of other clusters based on the gene expression in different treatment conditions and at different time points was not possible. t-SNE visualization of only those genes differentially expressed between the two treatments illustrates the lack of cluster resolution based on treatment or sampling time (Fig 4B).

### Shading results in downregulation of photosynthesis-, lysosome-, and carbon metabolism-related pathways

Pairwise comparisons of all shaded samples against illuminated samples (all samples from incubation A, sampling timepoint E1 from incubation B; Fig 1), shaded samples against illuminated samples excluding timepoint E1, and shaded samples against illuminated samples at each of timepoints E2, E3, and E4, yielded 424, 80, 23, 20, and 18 differentially expressed genes (DEGs) respectively. These DEGs mapped to five main KEGG Orthology pathway families: transcription and translation, photosynthetic activity, cell metabolism, cell stress response, and endocytic activity (Fig 5). In all these pathway families, except for photosynthetic activity, the DEGs were downregulated in the samples that were shaded.

**Figure 5:**
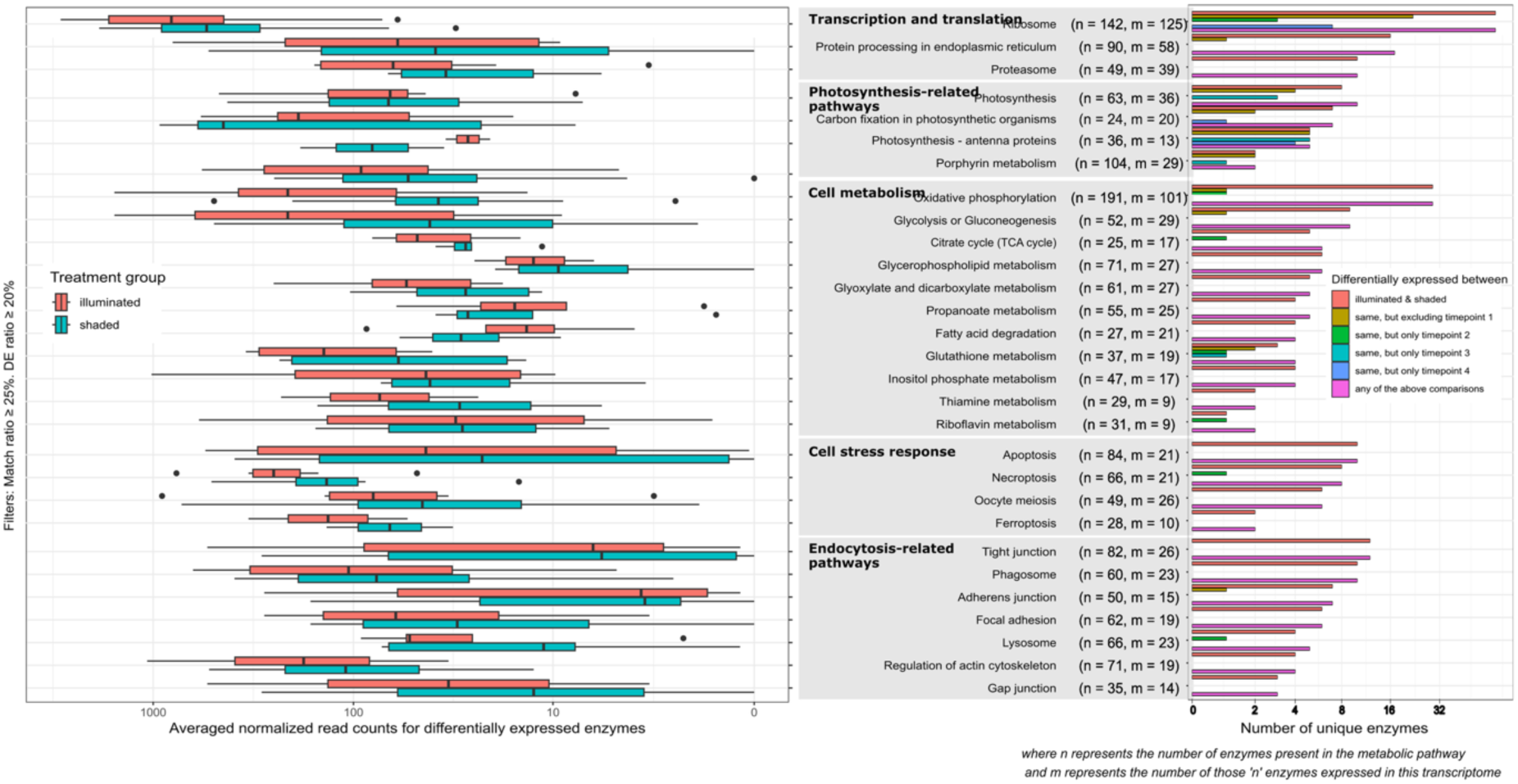
Metabolic mapping summary for *Rhodomonas*: Summary of KEGG pathways associated to the differentially expressed genes. For each pathway label, *n* represents the number of enzymes present in the pathway and *m* represents the number of those *n* enzymes recovered in the assembled *Rhodomonas* transcriptome. On the right side, bars represent number of differentially expressed genes for each pathway, coloured based on subsets of samples. Boxplots on the left side show the normalized counts of each of the differentially expressed genes averaged over all the samples in the given treatment group.

Pathways associated with transcription and translation had the highest number of DEGs: ribosome (70 DEGs), protein processing in endoplasmic reticulum (11 DEGs), and proteasome (10 DEGs). All the DEGs associated with this pathway family were downregulated in the shaded samples, which agrees with the overall gene expression pattern across the treatments (Fig 4A) and alludes to reduced transcriptional activity in the shaded samples.

Among the photosynthesis-related pathways – photosynthesis, carbon fixation in photosynthetic organisms, photosynthesis antenna proteins, and porphyrin metabolism – there were several genes that were upregulated in the shaded samples. Higher expression of protochlorophyllide reductase and protoporphyrin/coproporphyrin ferrochelatase suggest that the *Rhodomonas* cells attempt to increase the production of chlorophyll-*a* in response to the reducing light availability. This is further corroborated by upregulation of 4 of the 5 light-harvesting complex I proteins and 4 of the 6 photosystem II proteins. Downregulation of 5 of the 6 Calvin cycle proteins that were differentially expressed indicates lower availability of the ATP and NADPH from the light reactions due to the shading treatment.

Pathways associated with cellular stress responses included apoptosis, cellular senescence, necroptosis, meiosis, and ferroptosis, while pathways associated with endocytic activity included synthesis and activity of phagosomes and lysosomes, and regulation of actin cytoskeleton, tight junction, adherens junction, focal adhesion, and gap junction. Nearly all DEGs associated with these pathways had higher expression in illumination compared to shading. These observations, however limited, indicate that cells in the illuminated samples face high environmental stress but continue to feed, whereas the cells in the shaded samples show reduced lysosomal digestion and feeding.

The cell metabolism pathway family comprised the largest number of pathways with DEGs. These included oxidative phosphorylation, glycolysis, citrate cycle, and metabolism of glycerophospholipids, glyoxylates, propanoates, fatty acids, glutathione, inositol phosphate, thiamine, and riboflavin. Cells in the shaded treatment experienced a widespread downregulation of such cellular metabolism, confirmed by the downregulation of key enzymes in oxidative phosphorylation (all 5 complexes), glycolysis (pyruvate kinase), citrate cycle (citrate synthase), and *β* oxidation of fatty acids (long-chain acyl-CoA synthetase). Overexpression of isocitrate lyase (involved in the glyoxylate cycle) in the shaded samples indicates unavailability of simple sugars to satisfy cellular energy requirements. Lower expression of genes mediating the pathways of cellular metabolism and the overexpression of isocitrate lyase, together, imply a harsher energy landscape for the shaded samples.

## Discussion

We successfully sequenced the transcriptomes of 1,520 individual FACS-sorted microeukaryotes of environmental origin, belonging to 22 different families across 5 supergroups and 17 classes, without reference genomes or transcriptomes or continuous lab cultures. *De novo* partial transcriptomes were assembled for each of the 22 taxonomic groups and annotated using sequence-based and structure-based homology. For each of the 1,520 microeukaryote cells, we were able to accurately assign taxonomy, assemble *de novo* partial transcriptomes and analyse expression across two distinct light conditions over time. Additionally, we could identify their associated prokaryotes and map interdomain interactions. For Rhodomonas, the most abundant microeukaryote taxon in the study, we were able to map enough differentially expressed genes to KEGG metabolic maps to observe upregulation of light-harvesting complex proteins upon shading, suggesting an investment in limited light conditions, but downregulation of Calvin cycle enzymes. The subsequent scarcity of simple sugar substrates and downregulation of cellular metabolism in shaded samples was also observed.

We used Smart-seq3xpress^36^, as a service provided by Xpress Genomics, to prepare single-cell transcriptomic libraries for the microeukaryotes and sequence them. Since at least 100-150 taxonomically identical cells are needed to assemble a satisfactory transcriptome^17,34^, we limited the natural diversity of microeukaryotes to mixotrophs only by using chlorophyll *a* (for photoautotrophy) and LysoSensor Blue (for heterotrophy) fluorescence signals. However, we hypothesize that based on these signals alone, heterotrophs that are currently consuming/digesting photoautotrophs or have photoautotrophic bacteria attached to their surfaces are indistinguishable from mixotrophs. An overwhelming 87% of reads were UMI-end reads while only 13% were internal reads. On average, 33,263 unique UMIs were found per sample, but only 2643 mapped to the assembled transcriptomes. Although tagmentation enables recovery of variable 3’ ends from the same cDNA molecule^37^, the lack of internal reads limits the ability to assemble long transcripts, leading to incomplete transcriptomes and low UMI-end read mapping. However, this shows that broad functional characterization of microeukaryotes from complex natural communities is possible even with limited transcriptomes.

Expression of only about half of the OrthoDB eukaryota orthologous genes was recovered. It is possible that the incomplete recovery, indicated by missing conserved gene expression, may not be due to shortcomings of sequencing or transcriptome assembly, but due to aforementioned genes not being expressed at the time of sampling. Transcript accumulation curves (in Fig. 2E) for some of the transcriptomes indicate that the transcriptomes are incomplete and higher number of sampled cells would have yielded more complete transcriptomes. However, a comparison of the curves of *Kathablepharidida* and *Rhodomonas* indicates that in addition to number of sampled cells, completeness might be a function of sequencing depth, since 3.25M reads were on average recovered from *Kathablepharidida* cells, a number that is much higher than the study average. Following this line of argument and our previous study of *Ochromonas triangulata* transcriptomes^34^, we believe that 100M short reads, retrieved across multiple single cells, is sufficient to assemble a satisfactory transcriptome. The reads must be recovered from multiple single cells to infer cellular heterogeneity and because each individual microeukaryote cell only carries a small number of mRNA molecules^17^ and transcription in eukaryotes occurs in small bursts^38^.

Microeukaryote genomes and transcriptomes are understudied. Consequently, they are functionally uncharacterized, or their sequences are absent from annotation databases used to infer functionality. Even after using multiple databases (Uniprot-SwissProt^39^, AlphaFoldDB^40^, PDB^41^, KEGG^42^, Pfam^43^ etc) and sequence similarity as well as structure homology, less than 10% of the transcripts in the *Rhodomonas* transcriptome were functionally annotated. This will be the case for most microeukaryotes in natural environments, as they either lack references or cannot be cultured. Therefore, large-scale sequencing projects like P10K^14^ and MMETSP^13^ are needed to assemble reference genomes/transcriptomes and experimental studies targeting specific genes/proteins. Another potential explanation for the incomplete functional annotation is that many of the unannotated transcripts do not represent mature mRNAs and thus would need splicing, RNA editing etc. before function can be inferred from sequences. Molecular and computational methods to distinguish mature mRNAs need further development.

The most common bacterial families associated with the microeukaryotes were *Endomicrobiaceae* and *Leptolyngbyaceae*. Irrespective of treatments and sampling timepoints, the prokaryotic associates remained specific to the microeukaryotes sampled. Like macro-organisms, microeukaryotes have also been shown to accommodate communities of diverse co-existing prokaryotic associates^44–47^. Although microeukaryotes in the same natural microeukaryote population hosting different communities of prokaryotic symbionts have been observed^45^, our data shows that in the sampled freshwater community, microeukaryotes of the same species host diverse but similar prokaryotic associates and that these, mostly supergroup specific, associations are stable across differing light conditions and time, at least during the span of our experimental incubations. Common association of the morphologically and genetically diverse *Leptolyngbyaceae*^48^, known symbiote of mosses^49^ and sponges^50^, with the cryptophytes (*Rhodomonas*, *Kathablepharidida*, and *Cryptomonadales*) could indicate similar feeding behaviour across diverse cryptophytes. The association of *Endomicrobiaceae* intracellular endosymbionts with Alveolates (*Askenasia*) and Stramenopiles (*Pedinellales*, *Olisthodiscs*, *Cylindrotheca*, *Fragilariopsis*, and *Paraphysomonas*) could indicate a host specificity with the TSAR supergroup.

Following our previous reference-free study of *Ochromonas triangulata* (a marine mixotrophic microeukaryote)^34^ and the current reference-free and culture-free study of 22 different microeukaryotic taxa, we have set the stage for high-throughput single cell transcriptomics to be used to study the natural microeukaryotic diversity, in terms of population-level taxonomic identity, sub-population-level transcriptomic profiles, and individual-level prokaryotic associations. The number of individual single cell transcriptomes sequenced, and consequently the fraction of the natural microeukaryote diversity studied was limited by the throughput and cost of the sequencing strategy. With higher throughput techniques like droplet-based full-length transcript sequencing^51,52^ and split-and-pool barcoding^53,54^, we could scale 10X to 1000X the number of transcriptomes sequenced and decipher a larger fraction of the natural microeukaryote diversity.

## Online Methods

### Sample collection and conditioning

The sample was collected on 8th October 2024 from oligotrophic, clear water lake Långsjön (60° 1’ 58.5’’ N, 17° 34’ 47.1’’ E, TN = 0.63 mg L-1, TP = 9.8 μg L-1, Kd = 0.18 m-1). Water was collected at 1.5 m depth (T = 12°C) and filtered on-site through a 42 μm mesh-size nylon net into a clean collection container. Upon arrival to the laboratory (< 1 h from collection) the sample was transferred to a clean and autoclaved 5 L clear glass bottle and acclimatized for two days at 12°C and a photon flux of 80 μmol s-1 m-2 under a 14 h photoperiod.

### Experimental setup

Irradiance was manipulated in a one factor design, resulting in two treatments; (1) a control treatment, kept in acclimation conditions, and (2) a shading treatment, in which the light regime was reduced to 7.5% of acclimation irradiance. Both treatments were unreplicated. Initially, the sample was split into two 2 L glass bottles and incubated for 16 h prior to the start of the experimental phase (which included the full dark phase of the light cycle). At the start, 250 mL were sampled from each bottle for community metrics and single cell transcriptomics (E1), and immediately following E1, one incubation bottle was covered with two layers of 40 den black elastane stockings (shading treatment). Subsequently, identical samples were taken from each treatment after 6 h (E2), 24 h (E3) and 48 h (E4) after shading. Incubation bottles were sampled without refresh (i.e. adding sterile lake water to the incubation container in equal volume as the volume sampled), as this was observed to introduce stochastic variability in a pilot experiment (Supplementary Fig 1).

### Cell abundance and cytometric fingerprinting

All flow cytometry analyses were performed on a CytoFLEX S flow cytometer (Beckman Coulter, Brea, USA). Small volumes (20-30 mL) were sampled from the incubation bottles and were subsequently fractionated by filtering through a 70 μm mesh-size (fraction 1) and 1.2 μm (fraction 2) nylon cell strainers, and 0.1 μm pore-size (negative fraction) SUPOR membrane filters. SYBR Green II was used as fluorescent marker for cell enumeration (FL1, EX 488 EM 525/40 nm). Eukaryotes and prokaryotes were measured separately under different cytometer parameters. For eukaryotes, forward light scatter was used as trigger and cell abundance was estimated by subtracting the volumetric event count of fraction 2 in the FL1 channel to the same variable of fraction 1, both under the same acquisition settings. For prokaryotes, FL1 signal was used as trigger and cell abundance was approximated to the volumetric event count of fraction 2. In both cases, the negative fraction was used to establish the positivity threshold and set up the counting gates using side light scatter and FL1 signal.

Small aliquots (1-2 mL) were retrieved in triplicates from fraction 1 for cytometric fingerprinting. These aliquots were stained with SYBR Green II and LysoSensor Blue DND-147 (FL5, EM 405 EX 450/45) as a proxy for feeding vacuole, and cytometric data was recorded in the forward and side light scatter channels (FSC and SSC, respectively), FL1, FL5, FL2 (phycoerythrin autofluorescence, EX 488 EM 585/42) and FL3 (chlorophyll autofluorescence, EX 488 EM 690/50) channels.

### Single cell sorting and library preparation

Cell sorting was performed with a MoFlo Astrios EQ (Beckman Coulter) with active UV (355 nm), blue (488 nm) and red (640 nm) lasers. Sorting gates were drawn around putative mixotrophic eukaryotes based on events simultaneously positive in three channels: SYBR Green II (FL17, EX 488 EM 530/40), chlorophyll autofluorescence (FL23, EM 640 EX 671/30 nm) and LysoSensor Blue (FL7, EX 355 EM 448/59). Single cells were sorted onto 384-well plates containing lysis buffer (<500 nL) and 3.3 μL of a silicone overlay liquid (pre-loaded plates provided by Xpress Genomics). Plates were pulse-centrifuged immediately after sorting and stored at -80 °C until library preparation. Libraries were prepared by Xpress Genomics AB (Stockholm, Sweden) following the Smart-seq3xpress methodology. Sequencing was performed internally at Xpress Genomics on a DNBSEQ-G400 sequencer (MGI Tech, Shenzhen, China) in pair-end 150 bp configuration.

### Analysis of community trajectory using cytometric fingerprinting

Flow cytometry data was processed in R using the flowCore package (version 2.20.0, R version 4.4.2). For each sample and replicate, maximum signal intensity values (i.e. the height of the pulse) for all events in the FSC, SSC, FL1, FL2, FL3 and FL5 channels were recorded and mapped to a six-dimensional grid. The grid, binned on a uniform log 10 scale (10^2^-10^3^, 10^3^-10^4^, 10^4^-10^5^, 10^5^-10^6^, 10^6^-10^7^) along the six channels, contained 15,625 grid cells. The number of events in each grid cell were then counted and a distance matrix based on Bray-Curtis dissimilarity was computed for all samples and replicates. Only non-zero cells (i.e. grid cells that contain at least one event across all replicates) were considered to compute the matrix. Non-parametric multidimensional scaling was performed on the distance matrix using the metaMDS function of the vegan R package (version 2.6.10) using default parameters.

### Quality Check

Raw read sequences from Xpress-seq were quality filtered by Xpress Genomics AB (Stockholm, Sweden) to remove reads where barcode with more than 5 bases under Phred 20 or UMIs with more than 4 bases under Phred 20. Barcodes were error-corrected with 1 Hamming Distance. Barcodes and UMIs were extracted and returned to us as BAM tags in a BAM file containing sequences from all 1584 well plates. A custom python script was used to add the UMIs to the sequence headers and remove UMIs from sequences where multiple copies of the UMI were found at the beginning of the forward sequences. The BAM file was then split into BAM files for individual cells using the cell barcode encoded in the BC BAM tag using ‘bamtools split’ (version 2.5.2)^55^. Each BAM file was then split into two gzipped fastq files containing forward and reverse reads using ‘samtools fastq’ (version 1.21) ^56^. Nextera Transposase sequence adapter “CTGTCTCTTATA” were removed from the reads using Trim Galore^57^ (version 0.6.10) and Cutadapt^58^ (version 4.9). Trimmed reads were then filtered using Kraken2^59^ (version 2.1.2) to identify and separate bacterial and human contaminant sequences, with the standard database containing NCBI taxonomic information, as well as the complete genomes in RefSeq for the bacterial, archaeal, and viral domains, along with the human genome and a collection of known vectors (UniVec_Core).

### Ribosomal RNA read isolation and taxonomic assignment

RiboDetector^60^ (version 0.3.1) was used to separate ribosomal RNA reads from the rest of the raw reads. rRNA reads were used for taxonomic assignment of the microeukaryotes as well as their associated prokaryotes, using the tool vsearch^61^ (versions 2.22.1 and 2.29.2). Trimmed read pairs with a minimum merged sequence length of 150 were merged using the ‘vsearch --fastq_mergepairs’ command (version 2.22.1). The resulting merged sequences were dereplicated using the ‘vsearch -- fastx_uniques’ command (version 2.22.1) and written into a fasta file, while discarding sequences with a post-dereplication abundance value smaller than 10. The abundance of each dereplicated sequence in each of the samples was recorded using the ‘-- tabbedout’ option. The unique merged sequences were classified and assigned taxonomy using the Sintax algorithm^62^ implemented in the ‘vsearch --sintax’ command (version 2.29.2), and reported in a tab-separated text file using the ‘--tabbedout’ option. Version 2.29.2 was used for Sintax in order to use a random draw to break ties between sequences with equally many kmer matches as implemented in the ‘-- sintax_random’ option. A bootstrap support cutoff of 50% was enforced on the assigned taxonomy using a custom python script. Eukaryotic taxonomy was assigned to the sequences by using Sintax against the Protist Ribosomal Reference database (PR^2^) ^63^. The DADA2 fasta file of the PR^2^ version 5.0.0 release was modified using a custom python script to comply with the vsearch sintax input header requirements. The microeukaryote taxonomic assignment was then done by accounting for the highest abundance of annotated sequences in each sample and manual curation. Prokaryotic taxonomy was assigned to the sequences by using Sintax against the SILVA ribosomal RNA gene database^64^ (version 138.2). The non-redundant (NR, full reference dereplicated using a 99% identity criterion) rRNA small-subunit reference fasta database of the SILVA version 138.2 release was modified using a custom python script to comply with the vsearch sintax input header requirements. The prokaryote taxonomic assignments were then filtered to remove annotations corresponding to common laboratory-associated and human-associated contaminants^65^.

### *De novo* transcriptome assembly

Quality-filtered, adapter-trimmed, and contaminant-filtered reads from the same taxonomic group (usually genus, Supplementary table 1, 2) were pooled together and assembled into de novo partial reference transcriptomes using Trinity^66^ (version 2.15.2). In silico read normalization was performed to remove excess reads beyond 200x coverage with k-mers of length 25bp.

The quality of the assembled transcriptomes was assessed using multiple tools and metrics. The read representation of the assemblies was determined by aligning the reads to the assembled transcriptome using bowtie2^67^ (version 2.5.3) and counting the number of properly aligned read pairs as well as the improper or orphan read alignments. The completeness of the transcriptome assemblies was evaluated using compleasm^68^ (version 0.2.6) by aligning the assembled transcripts against the eukaryota BUSCO lineage orthologs^69,70^ and if available against more specific BUSCO lineage ortholog datasets (alveolata, ascomycota, chlorophyta, stramenopiles). The number of assembled transcripts that are full-length or nearly full-length were examined by blastx^71^ (version 2.15.0) searching them against the UniProt Swiss-Prot database^39^ with an e-value cutoff of 1e-10, and quantifying the percent of the target being aligned to by the best matching assembled transcript. Finally, the N50 transcript contig lengths as well as the ExN50 (where the N50 statistic is limited to the genes that represent x% of the total normalized expression) were calculated using perl scripts from the Trinity utilites^66^.

### Transcriptome annotation

The longest open reading frames in the *de novo* transcriptome contigs assembled by Trinity were using TransDecoder^72^ (version 5.7.1) with the ‘TransDecoder.LongOrfs’ command. Only open reading frames with at least 100 amino acids were retained. Since we weren’t interested in isoforms and gene variants, these sequences were then clustered at 90% global sequence identity with a word size of 5, using CD-HIT^73^ (version 4.8.1). The resulting non-redundant transcriptomes were annotated using different methods and tools and databases as described below. These were then homogenized and manually curated using R downstream.

#### Sequence homology

Trinotate^74^ (version 4.0.2) was used to automate the sequence homology based transcript nucleotide as well as open reading frame peptide sequences. Diamond blastx (version 2.1.10) was used to do a translated protein search of the entire assembled nucleotide transcript against the UniProt Swiss-Prot database^39,75^. Diamond blastp (version 2.1.10) was used to do a protein search of the non-redundant open reading frame peptide sequences against the UniProt Swiss-Prot database^39,75^. Protein domain identification was performed using HMMER (version 3.4) against the Pfam database^43,76^. Signal peptides were detected using a machine learning model called SignalP^77^ (version 6.0). The transmembrane protein domains were predicted using a deep learning model called DeepTMHMM^78^ (version 1.0.42). RNA homologs were identified using a covariance model based tool called Infernal^79^ (version 1.1.5). Functional annotation using precomputed orthologous groups was done using EggNOG-mapper^80^ (version 2.1.9). Multiple of the above methods were used to additionally include mapping to KEGG and GO databases.

The rRNA transcripts, identified by Infernal, using vsearch SINTAX, were searched against the Silva SSU & LSU databases (version 138.2) for bacterial reads, and PR^2^ SSU (version 5.0.0) and EUKARYOME LSU (version 1.9.4) databases for eukaryotic reads^63,64,81^ for confirmation of the rRNA read based taxonomy assignment.

#### Structure prediction and homology

ESMFold’s structure prediction model version 1 was used to predict atomic-level protein structures from the translated peptide sequences of the transcript open reading frames^82^. The model was downloaded from colabfold’s server and run locally using PyTorch version 1.12 in Python 3.9. Protein structures with predicted Local Distance Difference Test (pLDDT) score, a per-residue measure of local confidence, of less than 0.5 were filtered out. Structures with pLDDT of up to 0.7, up to 0.9, and over 0.9, were considered to be of good, high, and very qualities respectively.

Foldseek was then used to align these predicted structures against the Alphafold Swiss-Prot (version 4) which contains all the predicted structures of the proteins in the Swiss-Prot database as well as the Protein Data Bank’s 3D structures(dated 01.01.2024)^40,41,83^. Foldseek was run with the default sensitivity of 9.5 and 3Di+AA Gotoh-Smith-Waterman local alignment. Protein structure based function prediction was performed using a Graph Convolutional Network tool called DeepFRI^84^. For each protein structure, DeepFRI was used to predict the gene ontologies of biological process, molecular function, cell compartment, and also the enzyme commission classification.

#### Homogenizing annotation

For a given transcript in a given transcriptome, the protein identity was annotated in the following hierarchy: 1) Protein sequence (non-redundant open reading frame) homology search against UniProt Swiss-Prot, 2) Protein structure homology search against AlphaFoldDB Swiss-Prot, 3) Transcript sequence’s translated protein homology search against UniProt Swiss-Prot, 4) Protein structure homology search against the Protein Data Bank 3D structures, 5) Hidden Markov Model-based protein domain identification against PFam database, 6) EggNOG protein ortholog.

For the Gene Ontology, each transcript was annotated in the following ways: 1) Gene Ontology via UniProt ID obtained above (using the QuickGO API from the Protti R package), 2) Protein structure based Gene Ontology prediction (using DeepFRI), 3) EggNOG-mapper Gene Ontology assignment, 4) Gene Ontology mapping from the UniProt Swiss-Prot protein sequence homology search, 5) Gene Ontology mapping from the UniProt Swiss-Prot transcript sequence’s translated protein homology search. Gene Ontology annotations from each of the above ways were combined and the unique annotations per transcript were kept.

For the KEGG Orthologs, annotation derived from the UniProt IDs (via UniProt’s ID Mapping API and KEGG DBGET API) assigned above were preferred over EggNOG KEGG Ortholog mapping and those via Trinonate’s homology searches and KEGG database mappings.

### Read mapping and Unique Molecular Identifiers (UMIs)

The ‘align_and_estimate_abundance.pl’ perl script from the Trinity (version 2.15.2) utilities was used to map the quality-filtered, adapter-trimmed, and contaminant-filtered reads to the non-redundant (90% identity clustered) coding sequences for each of the transcriptomes. The perl script used Bowtie^85^ (version 1.2.3) for alignment and RSEM^86^ (version 1.3.3) for abundance estimation, with a maximum insert size of 800. The ‘abundance_estimates_to_matrix.pl’ perl script from the Trinity (version 2.15.2) utilities was used to tabulate the individually normalized counts (transcripts per million or TPM) for each different transcriptome and perform cross-sample normalization within each transcriptome using a weighted trimmed mean of the log expression ratios (trimmed mean of M values or TMM).

Then, internal reads were filtered out from the alignment. The UMIs were deduplicated in the alignment file using UMI-tools’ ‘umi_tools dedup’ command^87^ (version 1.1.6). The deduplicated UMI-containing read alignments were then, as above, used for abundance estimation using RSEM, TPM-normalized individually, and TMM-normalized across samples within each transcriptome. Cells with fewer than 100 distinct UMIs were discarded from downstream analyses.

### Transcript accumulation and coverage

Transcript accumulation and transcript coverage curves were generated by performing rarefaction and extrapolation using the iNEXT R package (version 3.0.1) ^88^. As curves for species accumulation and species coverage are generated in species diversity studies, transcript accumulation and transcript coverage curves were generated here, using cell identity in place of sampling site. Hill number (with q = 0) or species richness was used. For transcript accumulation curves to be comparable across transcriptomes, only the most conserved eukaryotic genes were sampled using compleasm (version 0.2.6) to map the transcripts against the eukaryota odb10 gene set ^68,70^.

### Gene expression clustering and visualization

Two-dimensional visualization of the genes commonly expressed among the chosen transcriptomes was made with exact t-distributed Stochastic Neighbour Embedding after reducing the initial number of dimensions to 50 with a truncated PCA and default parameters (momentum = 0.5, final momentum = 0.8, learning rate = 200, exaggeration factor = 12) and mean and standard deviation scaling. An R wrapper for Van der Maaten’s Barnes-Hut implementation of t-SNE in C++ was used (https://github.com/jkrijthe/Rtsne).

Gene expression was clustered using either shared nearest-neighbour (SNN) graph (with 13 nearest neighbours and Walktrap community finding algorithm) or hierarchical clustering (with Ward’s clustering criterion for agglomeration and Bray-Curtis dissimilarity, 1 – Pearson correlation, or cosine dissimilarity as the distance metrics). The hierarchical trees were cut at half the maximum height of the respective trees. Goodness of fit of the clusters was evaluated with pairwise Rand indices.

### Differential expression and metabolic mapping

The differential expression was estimated by counting a consensus of five different tools: ALDEx2, ANCOM-BC, MaAsLin2, LinDa, and DESeq2, because they use different strategies for transforming the data and correcting the bias due to the nature of the compositional data^89–94^. Only those genes found to be differentially expressed by at least two of the above five tools (with an adjusted p-value < 0.05) were considered to be differentially expressed. A 5% prevalence filter with a non-zero detection threshold was applied prior to tests for differential expression to remove rare features that may be artefacts.

Gene expression, in the form of KEGG Orthologs, was mapped to KEGG (Kyoto Encyclopedia of Genes and Genomes) Orthology pathways using the pathview R package (version 1.44.0)^42,95^. Only those pathways where at least 25% of proteins in the pathway were recovered in our transcriptome and at least 10% of these matched proteins were differentially expressed were retained for manual inspection.

Gene set enrichment analysis was performed using fgsea (version 1.34.0)^96^ with the Gene Ontology terms as the gene sets and the list of differentially expressed genes as the pre-ranked list and 10,000 permutations.

### Associated prokaryome analysis

The following analyses was done in R (version 4.4.1). As described above, rRNA reads were separated using RiboDetector, and searched for against the SILVA database using vsearch and SINTAX^60–62,64^. A bootstrap cutoff of 50% was applied. Resulting read counts were grouped per sample per species and summed together. The prokaryote taxonomic assignments were then filtered to remove annotations corresponding to common laboratory-associated and human-associated contaminants^65^. Data was formatted into a tree summarized experiment class of the mia package^97^ (version 1.17.3) and agglomerated at different taxonomic levels for visualization. The per-cell taxonomy and counts, along with some metadata, are reported in supplementary table 5. The counts were transformed to relative abundance for some downstream analyses.

Rarefaction curves were estimated using the ‘rarecurvè function from the vegan package (version 2.6-10), with a subsample size equal to the minimum of counts for each sample and a step size of 100.

Constrained metric scaling and redundancy analysis was done with a simple additive model to test how well the variance in the relative abundance was explained by the microeukaryote associate, the treatment, and the sampling timepoint, using the ‘capscalè function from the vegan package. A permutational test for the capscale model was done with Bray-Curtis dissimilarities with 999 permutations using the ‘anova.ccà function from the vegan package. The relative abundance data was agglomerated to the family taxonomic level for the above analyses.

The differential abundance was estimated at the family taxonomic level by counting a consensus of five different tools: ALDEx2, ANCOM-BC, MaAsLin2, LinDa, and DESeq2, because they use different strategies for transforming the data and correcting the bias due to the nature of the compositional data^89–93,98^. Prokaryotes were tested for differential abundance between each pair of the twelve most abundant microeukaryotes, and between the illuminated and shaded treatments for each of those twelve. Only those prokaryotes found to be differentially abundant (adjusted p-value < 0.05) by at least three of the above five tools were considered to be differentially abundant.

## Extended data

1. Supplementary table 1: Manually curated microeukaryote taxonomic identity
2. Supplementary table 2: Table with distribution of samples of the different transcriptomes across each treatment and sampling timepoint
3. Supplementary table 3: List of differentially expressed genes between the treatments for *Rhodomonas*. Column 1 indicates the differentially expressed gene. Columns 2 provides the number of packages that agree on the significance of differential expression. Columns 4-8 indicate individual packages’ inference of differential expression. Columns 9-56 provide detailed outputs from individual packages.
4. Supplementary table 4: List of differentially abundant families of prokaryotes associated with the different microeukaryotes. Column 1 indicates which between which two microeukaryotes the bacterial family in column 2 was differentially associated. Columns 3 provides the number of differential abundance packages that agree on the significance of differential abundance. Columns 4-8 indicate individual packages’ inference of differential abundance. Columns 9-56 provide detailed outputs from individual packages.
5. Supplementary table 5: List of prokaryotes, agglomerated to the family taxonomic level, found associated with each microeukaryote. Counts and cluster affiliation is also presented.
6. Supplementary figure 1: Non-parametric multidimensional scaling (NMDS) based on a cytometric fingerprinting analysis shows that replenishing an incubated lake sample with equivalent volume of sterile lake water after sampling—R+ treatment— results in larger stochastic variability than when the incubate is sampled but not replenished—R-treatment. Briefly, a lake water sample from lake Erken was collected and split into 9 x 1 L fractions. These fractions were all incubated simultaneously under identical regimes during 6 days but were subject to 3 different sampling schemes: 3 fractions were sampled daily (100 mL) and refreshed with an equivalent volume of sterilized natural lake water (R+ treatment), 3 fractions were sampled daily (100 mL) but not refreshed (R-treatment) and 3 fractions were not sampled at all (C-treatment). Flow cytometry data was collected at day 6 and further analysed as described the methods section.

## Acknowledgements

This work was partly financed by the Research Council of Finland (grants 323663 and 336475). The authors wish to acknowledge CSC – IT Center for Science, Finland, for computational resources.

## Data Availability Statement

Raw sequencing data will be uploaded to NCBI SRA and made available publicly upon the publication of this manuscript. Assembled and annotated transcriptomes will be made publicly available via github or the P10K database upon the publication of this manuscript.

## References

1. Bar-On YM, Phillips R, Milo R. The biomass distribution on Earth. Proceedings of the National Academy of Sciences. 2018;115(25):6506–6511. doi:10.1073/pnas.1711842115

2. Burki F, Roger AJ, Brown MW, Simpson AGB. The New Tree of Eukaryotes. Trends in Ecology & Evolution. 2020;35(1):43–55. doi:10.1016/j.tree.2019.08.008

3. Keeling PJ, Burki F. Progress towards the Tree of Eukaryotes. Current Biology. 2019;29(16):R808–R817. doi:10.1016/j.cub.2019.07.031

4. Mitra A, Caron DA, Faure E, et al. The Mixoplankton Database (MDB): Diversity of photo-phago-trophic plankton in form, function, and distribution across the global ocean. Journal of Eukaryotic Microbiology. 2023;70(4):e12972. doi:10.1111/jeu.12972

5. Simpson AGB, Slamovits CH, Archibald JM. Protist Diversity and Eukaryote Phylogeny. In: Archibald JM, Simpson AGB, Slamovits CH, eds. Handbook of the Protists. Springer International Publishing; 2017:1–21. doi:10.1007/978-3-319-28149-0_45

6. Smith DR, Keeling PJ. Protists and the Wild, Wild West of Gene Expression: New Frontiers, Lawlessness, and Misfits. Annual Review of Microbiology. 2016;70(2016):161–178. doi:10.1146/annurev-micro-102215-095448

7. del Campo J, Sieracki ME, Molestina R, Keeling P, Massana R, Ruiz-Trillo I. The others: our biased perspective of eukaryotic genomes. Trends Ecol Evol. 2014;29(5):252–259. doi:10.1016/j.tree.2014.03.006

8. Caron DA, Alexander H, Allen AE, et al. Probing the evolution, ecology and physiology of marine protists using transcriptomics. Nat Rev Microbiol. 2017;15(1):6–20. doi:10.1038/nrmicro.2016.160

9. Altenhoff AM, Glover NM, Dessimoz C. Inferring Orthology and Paralogy. In: Anisimova M, ed. Evolutionary Genomics. Vol 1910. Methods in Molecular Biology. Springer New York; 2019:149–175. doi:10.1007/978-1-4939-9074-0_5

10. Shazib SUA, Ahsan R, Leleu M, McManus GB, Katz LA, Santoferrara LF. Phylogenomic workflow for uncultivable microbial eukaryotes using single-cell RNA sequencing − A case study with planktonic ciliates (Ciliophora, Oligotrichea). Molecular Phylogenetics and Evolution. 2025;204:108239. doi:10.1016/j.ympev.2024.108239

11. Hackett JD, Anderson DM, Erdner DL, Bhattacharya D. Dinoflagellates: a remarkable evolutionary experiment. American Journal of Botany. 2004;91(10):1523–1534. doi:10.3732/ajb.91.10.1523

12. Amin SA, Hmelo LR, van Tol HM, et al. Interaction and signalling between a cosmopolitan phytoplankton and associated bacteria. Nature. 2015;522(7554):98–101. doi:10.1038/nature14488

13. Keeling PJ, Burki F, Wilcox HM, et al. The Marine Microbial Eukaryote Transcriptome Sequencing Project (MMETSP): Illuminating the Functional Diversity of Eukaryotic Life in the Oceans through Transcriptome Sequencing. PLOS Biology. 2014;12(6):e1001889. doi:10.1371/journal.pbio.1001889

14. Gao X, Chen K, Xiong J, et al. The P10K database: a data portal for the protist 10 000 genomes project. Nucleic Acids Research. 2024;52(D1):D747–D755. doi:10.1093/nar/gkad992

15. Pawlowski J, Audic S, Adl S, et al. CBOL Protist Working Group: Barcoding Eukaryotic Richness beyond the Animal, Plant, and Fungal Kingdoms. PLoS Biol. 2012;10(11):e1001419. doi:10.1371/journal.pbio.1001419

16. Kolisko M, Boscaro V, Burki F, Lynn DH, Keeling PJ. Single-cell transcriptomics for microbial eukaryotes. Current Biology. 2014;24(22):R1081–R1082. doi:10.1016/j.cub.2014.10.026

17. Liu Z, Hu SK, Campbell V, Tatters AO, Heidelberg KB, Caron DA. Single-cell transcriptomics of small microbial eukaryotes: limitations and potential. The ISME Journal. 2017;11(5):1282–1285. doi:10.1038/ismej.2016.190

18. Onsbring H, Tice AK, Barton BT, Brown MW, Ettema TJG. An efficient single-cell transcriptomics workflow for microbial eukaryotes benchmarked on Giardia intestinalis cells. BMC Genomics. 2020;21(1):1–9. doi:10.1186/s12864-020-06858-7

19. Jiang Y, Chen X, Wang C, et al. Genes and proteins expressed at different life cycle stages in the model protist Euplotes vannus revealed by both transcriptomic and proteomic approaches. Sci China Life Sci. 2025;68(1):232–248. doi:10.1007/s11427-023-2605-9

20. Reid AJ, Talman AM, Bennett HM, et al. Single-cell RNA-seq reveals hidden transcriptional variation in malaria parasites. Krzych U, ed. eLife. 2018;7:e33105. doi:10.7554/eLife.33105

21. Howick VM, Russell AJC, Andrews T, et al. The Malaria Cell Atlas: Single parasite transcriptomes across the complete Plasmodium life cycle. Science. 2019;365(6455):eaaw2619. doi:10.1126/science.aaw2619

22. Nadal-Ribelles M, Islam S, Wei W, et al. Sensitive high-throughput single-cell RNA-seq reveals within-clonal transcript correlations in yeast populations. Nat Microbiol. 2019;4(4):683–692. doi:10.1038/s41564-018-0346-9

23. Saint M, Bertaux F, Tang W, et al. Single-cell imaging and RNA sequencing reveal patterns of gene expression heterogeneity during fission yeast growth and adaptation. Nat Microbiol. 2019;4(3):480–491. doi:10.1038/s41564-018-0330-4

24. Nishimura Y, Otagiri M, Yuki M, et al. Division of functional roles for termite gut protists revealed by single-cell transcriptomes. ISME J. 2020;14(10):2449–2460. doi:10.1038/s41396-020-0698-z

25. Juéry C, Auladell A, Füssy Z, et al. Transportome remodeling of a symbiotic microalga inside a planktonic host. ISME J. 2024;18(1):wrae239. doi:10.1093/ismejo/wrae239

26. Zhou W, Zhao W, Yang S, et al. Single-cell transcriptome profiles and E−64 inhibitor data reveal the essential role of cysteine proteases in the ontogeny of *Ichthyophthirius multifiliis*. Fish & Shellfish Immunology. 2024;154:109979. doi:10.1016/j.fsi.2024.109979

27. Yan Y, Maurer-Alcalá XX, Knight R, Kosakovsky Pond SL, Katz LA. Single-Cell Transcriptomics Reveal a Correlation between Genome Architecture and Gene Family Evolution in Ciliates. mBio. 2019;10(6):10.1128/mbio.02524-19. doi:10.1128/mbio.02524-19

28. Onsbring H, Jamy M, Ettema TJG. RNA Sequencing of Stentor Cell Fragments Reveals Transcriptional Changes during Cellular Regeneration. Current Biology. 2018;28(8):1281–1288.e3. doi:10.1016/j.cub.2018.02.055

29. Cooney EC, Holt CC, Hehenberger E, Adams JA, Leander BS, Keeling PJ. Investigation of heterotrophs reveals new insights in dinoflagellate evolution. Molecular Phylogenetics and Evolution. 2024;196:108086. doi:10.1016/j.ympev.2024.108086

30. Galindo LJ, Mathur V, Frost H, Torruella G, Richards TA, Irwin NAT. Transcriptomics of Diphyllatea (CRuMs) from South Pacific crater lakes confirm new cryptic clades. Journal of Eukaryotic Microbiology. 2024;71(6):e13060. doi:10.1111/jeu.13060

31. Mtawali M, Cooney EC, Adams J, Jin J, Holt CC, Keeling PJ. Phylogenomic resolution of marine to freshwater dinoflagellate transitions. ISME J. 2025;19(1):wraf031. doi:10.1093/ismejo/wraf031

32. Ku C, Sebé-Pedrós A. Using single-cell transcriptomics to understand functional states and interactions in microbial eukaryotes. Philosophical Transactions of the Royal Society B: Biological Sciences. 2019;374(1786):20190098. doi:10.1098/rstb.2019.0098

33. Onsbring H, Tice AK, Barton BT, Brown MW, Ettema TJG. An efficient single-cell transcriptomics workflow for microbial eukaryotes benchmarked on Giardia intestinalis cells. BMC Genomics. 2020;21(1):1–9. doi:10.1186/s12864-020-06858-7

34. Jeevannavar A, Florenza J, Divne AM, Tamminen M, Bertilsson S. Cellular heterogeneity in metabolism and associated microbiome of a non-model phytoflagellate. The ISME Journal. 2025;19(1):wraf046. doi:10.1093/ismejo/wraf046

35. Grujčić V, Saarenpää S, Sundh J, et al. Towards high-throughput parallel imaging and single-cell transcriptomics of microbial eukaryotic plankton. PLOS ONE. 2024;19(1):e0296672. doi:10.1371/journal.pone.0296672

36. Hagemann-Jensen M, Ziegenhain C, Sandberg R. Scalable single-cell RNA sequencing from full transcripts with Smart-seq3xpress. Nat Biotechnol. 2022;40(10):1452–1457. doi:10.1038/s41587-022-01311-4

37. Hagemann-Jensen M, Ziegenhain C, Chen P, et al. Single-cell RNA counting at allele and isoform resolution using Smart-seq3. Nat Biotechnol. 2020;38(6):708–714. doi:10.1038/s41587-020-0497-0

38. Zenklusen D, Larson DR, Singer RH. Single-RNA counting reveals alternative modes of gene expression in yeast. Nat Struct Mol Biol. 2008;15(12):1263–1271. doi:10.1038/nsmb.1514

39. The UniProt Consortium. UniProt: the Universal Protein Knowledgebase in 2025. Nucleic Acids Research. 2025;53(D1):D609–D617. doi:10.1093/nar/gkae1010

40. Bordin N, Sillitoe I, Nallapareddy V, et al. AlphaFold2 reveals commonalities and novelties in protein structure space for 21 model organisms. Commun Biol. 2023;6(1):1–12. doi:10.1038/s42003-023-04488-9

41. Burley SK, Bhikadiya C, Bi C, et al. RCSB Protein Data Bank: powerful new tools for exploring 3D structures of biological macromolecules for basic and applied research and education in fundamental biology, biomedicine, biotechnology, bioengineering and energy sciences. Nucleic Acids Research. 2021;49(D1):D437–D451. doi:10.1093/nar/gkaa1038

42. Kanehisa M, Furumichi M, Sato Y, Matsuura Y, Ishiguro-Watanabe M. KEGG: biological systems database as a model of the real world. Nucleic Acids Research. 2025;53(D1):D672–D677. doi:10.1093/nar/gkae909

43. Finn RD, Mistry J, Tate J, et al. The Pfam protein families database. Nucleic Acids Research. 2010;38(suppl_1):D211–D222. doi:10.1093/nar/gkp985

44. Rossi A, Bellone A, Fokin SI, Boscaro V, Vannini C. Detecting Associations Between Ciliated Protists and Prokaryotes with Culture-Independent Single-Cell Microbiomics: a Proof-of-Concept Study. Microb Ecol. 2019;78(1):232–242. doi:10.1007/s00248-018-1279-9

45. Boscaro V, Manassero V, Keeling PJ, Vannini C. Single-cell Microbiomics Unveils Distribution and Patterns of Microbial Symbioses in the Natural Environment. Microb Ecol. 2023;85(1):307–316. doi:10.1007/s00248-021-01938-x

46. Zhang X, Bi L, Gentekaki E, Zhao J, Shen P, Zhang Q. Culture-Independent Single-Cell PacBio Sequencing Reveals Epibiotic Variovorax and Nucleus Associated Mycoplasma in the Microbiome of the Marine Benthic Protist Geleia sp. YT (Ciliophora, Karyorelictea). Microorganisms. 2023;11(6):1500. doi:10.3390/microorganisms11061500

47. Husnik F, Tashyreva D, Boscaro V, George EE, Lukeš J, Keeling PJ. Bacterial and archaeal symbioses with protists. Current Biology. 2021;31(13):R862–R877. doi:10.1016/j.cub.2021.05.049

48. Albertano P, Kováčik Ĺ. Is the genus Leptolyngbya (Cyanophyte) a homogeneous taxon? archiv_algolstud. 1995;75:37–51. doi:10.1127/algol_stud/75/1995/37

49. Lobakova ES, Zaytseva AA, Shibzukhova KA, Vasilieva SG, Butaeva GB, Gorelova OA. Cyanobacteria of moss symbioses from White Sea coast: ultrastructural characteristics and taxonomic diversity. Symbiosis. 2025;96(1):77–90. doi:10.1007/s13199-025-01062-1

50. Konstantinou D, Gerovasileiou V, Voultsiadou E, Gkelis S. Sponges-Cyanobacteria associations: Global diversity overview and new data from the Eastern Mediterranean. PLOS ONE. 2018;13(3):e0195001. doi:10.1371/journal.pone.0195001

51. Clark IC, Fontanez KM, Meltzer RH, et al. Microfluidics-free single-cell genomics with templated emulsification. Nat Biotechnol. 2023;41(11):1557–1566. doi:10.1038/s41587-023-01685-z

52. Sun X, Dadon SL, Ennis D, et al. Expanding single-cell toolbox with a cost-effective full-length total RNA droplet-based sequencing technology. bioRxiv. Preprint posted online February 27, 2025:2025.02.23.639726. doi:10.1101/2025.02.23.639726

53. Rosenberg AB, Roco CM, Muscat RA, et al. Single-cell profiling of the developing mouse brain and spinal cord with split-pool barcoding. Science. 2018;360(6385):176–182. doi:10.1126/science.aam8999

54. Martin BK, Qiu C, Nichols E, et al. Optimized single-nucleus transcriptional profiling by combinatorial indexing. Nat Protoc. 2023;18(1):188–207. doi:10.1038/s41596-022-00752-0

55. Barnett DW, Garrison EK, Quinlan AR, Strömberg MP, Marth GT. BamTools: a C++ API and toolkit for analyzing and managing BAM files. Bioinformatics. 2011;27(12):1691–1692. doi:10.1093/bioinformatics/btr174

56. Danecek P, Bonfield JK, Liddle J, et al. Twelve years of SAMtools and BCFtools. GigaScience. 2021;10(2):giab008. doi:10.1093/gigascience/giab008

57. Krueger F, James F, Ewels P, et al. FelixKrueger/TrimGalore: v0.6.10 -add default decompression path. Published online February 2, 2023. doi:10.5281/zenodo.7598955

58. Martin M. Cutadapt removes adapter sequences from high-throughput sequencing reads. EMBnet.journal. 2011;17(1):10–12. doi:10.14806/ej.17.1.200

59. Wood DE, Lu J, Langmead B. Improved metagenomic analysis with Kraken 2. Genome Biol. 2019;20(1):1–13. doi:10.1186/s13059-019-1891-0

60. Deng ZL, Münch PC, Mreches R, McHardy AC. Rapid and accurate identification of ribosomal RNA sequences via deep learning. Nucleic Acids Res. 2022;50(10):e60. doi:10.1093/nar/gkac112

61. Rognes T, Flouri T, Nichols B, Quince C, Mahé F. VSEARCH: a versatile open source tool for metagenomics. PeerJ. 2016;4:e2584. doi:10.7717/peerj.2584

62. Edgar RC. SINTAX: a simple non-Bayesian taxonomy classifier for 16S and ITS sequences. *bioRxiv*. Preprint posted online September 9, 2016:074161. doi:10.1101/074161

63. Guillou L, Bachar D, Audic S, et al. The Protist Ribosomal Reference database (PR2): a catalog of unicellular eukaryote Small Sub-Unit rRNA sequences with curated taxonomy. Nucleic Acids Research. 2012;41(D1):D597–D604. doi:10.1093/nar/gks1160

64. Quast C, Pruesse E, Yilmaz P, et al. The SILVA ribosomal RNA gene database project: improved data processing and web-based tools. Nucleic Acids Research. 2013;41(D1):D590–D596. doi:10.1093/nar/gks1219

65. Salter SJ, Cox MJ, Turek EM, et al. Reagent and laboratory contamination can critically impact sequence-based microbiome analyses. BMC Biol. 2014;12(1):1–12. doi:10.1186/s12915-014-0087-z

66. Grabherr MG, Haas BJ, Yassour M, et al. Full-length transcriptome assembly from RNA-Seq data without a reference genome. Nat Biotechnol. 2011;29(7):644–652. doi:10.1038/nbt.1883

67. Langmead B, Salzberg SL. Fast gapped-read alignment with Bowtie 2. Nat Methods. 2012;9(4):357–359. doi:10.1038/nmeth.1923

68. Huang N, Li H. compleasm: a faster and more accurate reimplementation of BUSCO. Bioinformatics. 2023;39(10):btad595. doi:10.1093/bioinformatics/btad595

69. Manni M, Berkeley MR, Seppey M, Simão FA, Zdobnov EM. BUSCO Update: Novel and Streamlined Workflows along with Broader and Deeper Phylogenetic Coverage for Scoring of Eukaryotic, Prokaryotic, and Viral Genomes. Molecular Biology and Evolution. 2021;38(10):4647–4654. doi:10.1093/molbev/msab199

70. Zdobnov EM, Kuznetsov D, Tegenfeldt F, Manni M, Berkeley M, Kriventseva EV. OrthoDB in 2020: evolutionary and functional annotations of orthologs. Nucleic Acids Research. 2021;49(D1):D389–D393. doi:10.1093/nar/gkaa1009

71. Camacho C, Coulouris G, Avagyan V, et al. BLAST+: architecture and applications. BMC Bioinformatics. 2009;10:421. doi:10.1186/1471-2105-10-421

72. Brian J. Haas. TransDecoder. Published online May 20, 2025. Accessed May 28, 2025. https://github.com/TransDecoder/TransDecoder

73. Fu L, Niu B, Zhu Z, Wu S, Li W. CD-HIT: accelerated for clustering the next-generation sequencing data. Bioinformatics. 2012;28(23):3150–3152. doi:10.1093/bioinformatics/bts565

74. Bryant DM, Johnson K, DiTommaso T, et al. A Tissue-Mapped Axolotl De Novo Transcriptome Enables Identification of Limb Regeneration Factors. Cell Reports. 2017;18(3):762–776. doi:10.1016/j.celrep.2016.12.063

75. Buchfink B, Reuter K, Drost HG. Sensitive protein alignments at tree-of-life scale using DIAMOND. Nat Methods. 2021;18(4):366–368. doi:10.1038/s41592-021-01101-x

76. Eddy SR. Accelerated Profile HMM Searches. PLOS Computational Biology. 2011;7(10):e1002195. doi:10.1371/journal.pcbi.1002195

77. Teufel F, Almagro Armenteros JJ, Johansen AR, et al. SignalP 6.0 predicts all five types of signal peptides using protein language models. Nat Biotechnol. 2022;40(7):1023–1025. doi:10.1038/s41587-021-01156-3

78. Hallgren J, Tsirigos KD, Pedersen MD, et al. DeepTMHMM predicts alpha and beta transmembrane proteins using deep neural networks. Bioinformatics. Preprint posted online April 10, 2022. doi:10.1101/2022.04.08.487609

79. Nawrocki EP, Eddy SR. Infernal 1.1: 100-fold faster RNA homology searches. Bioinformatics. 2013;29(22):2933–2935. doi:10.1093/bioinformatics/btt509

80. Cantalapiedra CP, Hernández-Plaza A, Letunic I, Bork P, Huerta-Cepas J. eggNOG-mapper v2: Functional Annotation, Orthology Assignments, and Domain Prediction at the Metagenomic Scale. Molecular Biology and Evolution. 2021;38(12):5825–5829. doi:10.1093/molbev/msab293

81. Tedersoo L, Hosseyni Moghaddam MS, Mikryukov V, et al. EUKARYOME: the rRNA gene reference database for identification of all eukaryotes. Database. 2024;2024:baae043. doi:10.1093/database/baae043

82. Lin Z, Akin H, Rao R, et al. Evolutionary-scale prediction of atomic-level protein structure with a language model. Science. 2023;379(6637):1123–1130. doi:10.1126/science.ade2574

83. van Kempen M, Kim SS, Tumescheit C, et al. Fast and accurate protein structure search with Foldseek. Nat Biotechnol. 2024;42(2):243–246. doi:10.1038/s41587-023-01773-0

84. Gligorijević V, Renfrew PD, Kosciolek T, et al. Structure-based protein function prediction using graph convolutional networks. Nat Commun. 2021;12(1):3168. doi:10.1038/s41467-021-23303-9

85. Langmead B, Trapnell C, Pop M, Salzberg SL. Ultrafast and memory-efficient alignment of short DNA sequences to the human genome. Genome Biol. 2009;10(3):1–10. doi:10.1186/gb-2009-10-3-r25

86. Li B, Dewey CN. RSEM: accurate transcript quantification from RNA-Seq data with or without a reference genome. BMC Bioinformatics. 2011;12(1):1–16. doi:10.1186/1471-2105-12-323

87. Smith TS, Heger A, Sudbery I. UMI-tools: Modelling sequencing errors in Unique Molecular Identifiers to improve quantification accuracy. Genome Res. Published online January 18, 2017:gr.209601.116. doi:10.1101/gr.209601.116

88. Hsieh TC, Ma KH, Chao A. iNEXT: an R package for rarefaction and extrapolation of species diversity (Hill numbers). Methods in Ecology and Evolution. 2016;7(12):1451–1456. doi:10.1111/2041-210X.12613

89. Fernandes AD, Reid JN, Macklaim JM, McMurrough TA, Edgell DR, Gloor GB. Unifying the analysis of high-throughput sequencing datasets: characterizing RNA-seq, 16S rRNA gene sequencing and selective growth experiments by compositional data analysis. Microbiome. 2014;2(1):1–13. doi:10.1186/2049-2618-2-15

90. Lin H, Peddada SD. Analysis of compositions of microbiomes with bias correction. Nat Commun. 2020;11(1):3514. doi:10.1038/s41467-020-17041-7

91. Mallick H, Rahnavard A, McIver LJ, et al. Multivariable association discovery in population-scale meta-omics studies. PLOS Computational Biology. 2021;17(11):e1009442. doi:10.1371/journal.pcbi.1009442

92. Zhou H, He K, Chen J, Zhang X. LinDA: linear models for differential abundance analysis of microbiome compositional data. Genome Biol. 2022;23(1):1–23. doi:10.1186/s13059-022-02655-5

93. Love MI, Huber W, Anders S. Moderated estimation of fold change and dispersion for RNA-seq data with DESeq2. Genome Biol. 2014;15(12):1–21. doi:10.1186/s13059-014-0550-8

94. Costa-Silva J, Domingues D, Lopes FM. RNA-Seq differential expression analysis: An extended review and a software tool. PLOS ONE. 2017;12(12):e0190152. doi:10.1371/journal.pone.0190152

95. Luo W, Brouwer C. Pathview: an R/Bioconductor package for pathway-based data integration and visualization. Bioinformatics. 2013;29(14):1830–1831. doi:10.1093/bioinformatics/btt285

96. Korotkevich G, Sukhov V, Budin N, Shpak B, Artyomov MN, Sergushichev A. Fast gene set enrichment analysis. bioRxiv. Published online 2021. doi:10.1101/060012

97. Tuomas Borman, Felix G.M. Ernst, Sudarshan A. Shetty, Leo Lahti. mia: Microbiome Analysis. doi:10.18129/B9.BIOC.MIA

98. Nearing JT, Douglas GM, Hayes MG, et al. Microbiome differential abundance methods produce different results across 38 datasets. Nat Commun. 2022;13(1):342. doi:10.1038/s41467-022-28034-z

