## Supplementary figures and images for "Culture-free single-cell transcriptomics shows short-term functional and associative stability in non-model microeukaryotes under shading stress"

### Supplementary figure 1

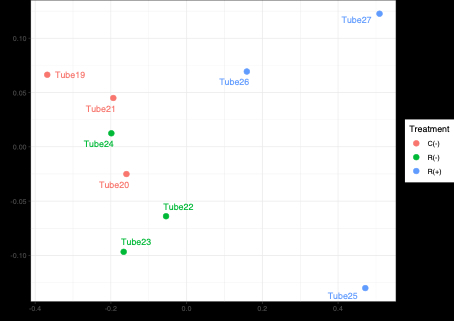
